# A dynamical motif comprising the interactions between antigens and CD8 T cells may underlie the outcomes of viral infections

**DOI:** 10.1101/540054

**Authors:** Subhasish Baral, Rustom Antia, Narendra M. Dixit

**Affiliations:** Department of Chemcial Engineering, Indian Institute of Science, Bangalore, India; Department of Biology, Emory University, Atlanta, USA; Centre for Biosystems Science and Engineering, Indian Institute of Science, Bangalore, India

## Abstract

Some viral infections culminate in very different outcomes in different individuals. They can be rapidly cleared in some, cause persistent infection in others, and mortality from immunopathology in yet others. The conventional view is that the different outcomes arise as a consequence of the complex interactions between a large number of different factors (virus, different immune cells and cytokines). Here, we identify a simple dynamical motif comprising the essential interactions between antigens and CD8 T cells and posit it as predominantly determining the outcomes. Antigen can activate CD8 T cells, which in turn can kill infected cells. Sustained antigen stimulation, however, can cause CD8 T cell exhaustion, compromising effector function. Using mathematical modelling, we show that the motif comprising these interactions recapitulates all the outcomes observed. The motif presents a new conceptual framework to understand the variable outcomes of infection. It also explains a number of confounding experimental observations, including the variation in outcomes with the viral inoculum size, the evolutionary advantage of exhaustion in preventing lethal pathology, the ability of NK cells to act as rheostats tuning outcomes, and the role of the innate immune response in the spontaneous clearance of hepatitis C. Interventions that modulate the interactions in the motif may present novel routes to clear persistent infections or limit immunopathology.

---

**V**iral infection can have different outcomes in different individuals. Three frequently observed outcomes are clearance after acute infection; long-term persistence; and host mortality due to immunopathology. With the hepatitis C virus (HCV), for instance, *∼*25% of the individuals infected clear the infection spontaneously, whereas the rest CD8+ chronically infected (1,2). Hepatitis B virus (HBV) causes persistent infection in neonates but is typically self-limiting in adults (3). The extent of pathology after infection can also vary; for example, respiratory syncytial virus (RSV) infection causes severe immunopathology in some children but not in others (4). Why can infection with a given virus have different outcomes in different individuals and what determines the outcome?

Mice models of lymphocytic choriomeningitis virus (LCMV) infection recapitulate many of these outcomes and consequently have been employed to better understand the factors that determine the outcomes. A seminal study, over 25 years ago, showed the effect of viral inoculum size on the outcome of LCMV infection (5): Infections initiated with a small virus inoculum were cleared, infections with intermediate inocula caused severe immunopathology leading to death, and infections with large inocula led to survival, albeit with the persistence of high titers of the virus (5, 6). Soon after, CD8 T cells were identified to be critical to the control of LCMV infections: CD8 T cell deficiency resulted in persistent infection of mice that otherwise would have rapidly cleared the virus (7). More recently, natural killer (NK) cells have been argued to act as rheostats tuning outcomes: NK cell depletion can change the outcome of infection with a high dose of LCMV from persistence of the virus with low levels of pathology to rapid host death (6). In other studies, regulatory T cells (8), factors contributing to CD8 T cell dysfunction (9, 10), type I interferon (IFN) signalling (11, 12), and, most recently, host genetics (13) have been argued to influence the outcomes of LCMV infection. A plethora of factors thus appears to be involved in determining the outcomes, and it is unclear how the interplay between these factors determines which outcome is reached.

Here, we identify a dynamical motif comprising the essential interactions of antigens and CD8 T cells and posit it as central to defining the outcomes of viral infections. Our reasoning follows from observations that suggest that the latter interactions may be both necessary and adequate to recapitulate the outcomes observed. Antigens exert competing influences on CD8 T cells. Antigen stimulation, via viral peptides presented by class I major histocompatibility complex (MHC-I) molecules on infected cells, activates CD8 T cells and triggers a response that can kill the infected cells (14, 15). Sustained stimulation, however, when a high level of antigen is present for an extended duration, can cause CD8 T cell dysfunction via exhaustion (5, 16–18). The relative strengths of these competing influences together with the strength of the CD8 T cell response appear to define the outcomes realized. While a strong CD8 T cell response is critical to viral clearance (7), persistent infections are typically associated with a dysfunctional CD8 T cell response (7, 14, 19). Furthermore, T cell exhaustion has been proposed to be an altered differentiation program to combat chronic infection while preventing immunopathology (20–22). The exhausted state is maintained by inhibitory molecules such as PD-1 (10) and can be partly reversed by blocking the inhibitory molecules or lowering antigen burden (9, 10, 23), which in turn can alter the disease state from persistence to clearance (24). We hypothesized, therefore, that the dynamical motif due to these underlying interactions governs the outcomes. Furthermore, we suggest that the other factors that influence the outcomes do so by modulating the interactions in the motif.

## Results

### The dynamical motif and its mathematical model

To test the hypothesis, we constructed the dynamical motif comprising the competing influences of antigen on CD8 T cells and the suppressive influence of CD8 T cells on antigen (Figure 1A). We developed the following minimal mathematical model to analyze the motif:

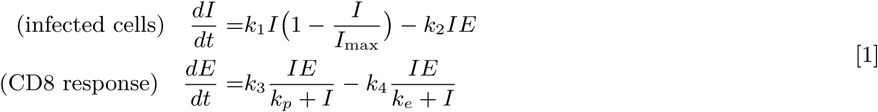

**Fig. 1.**
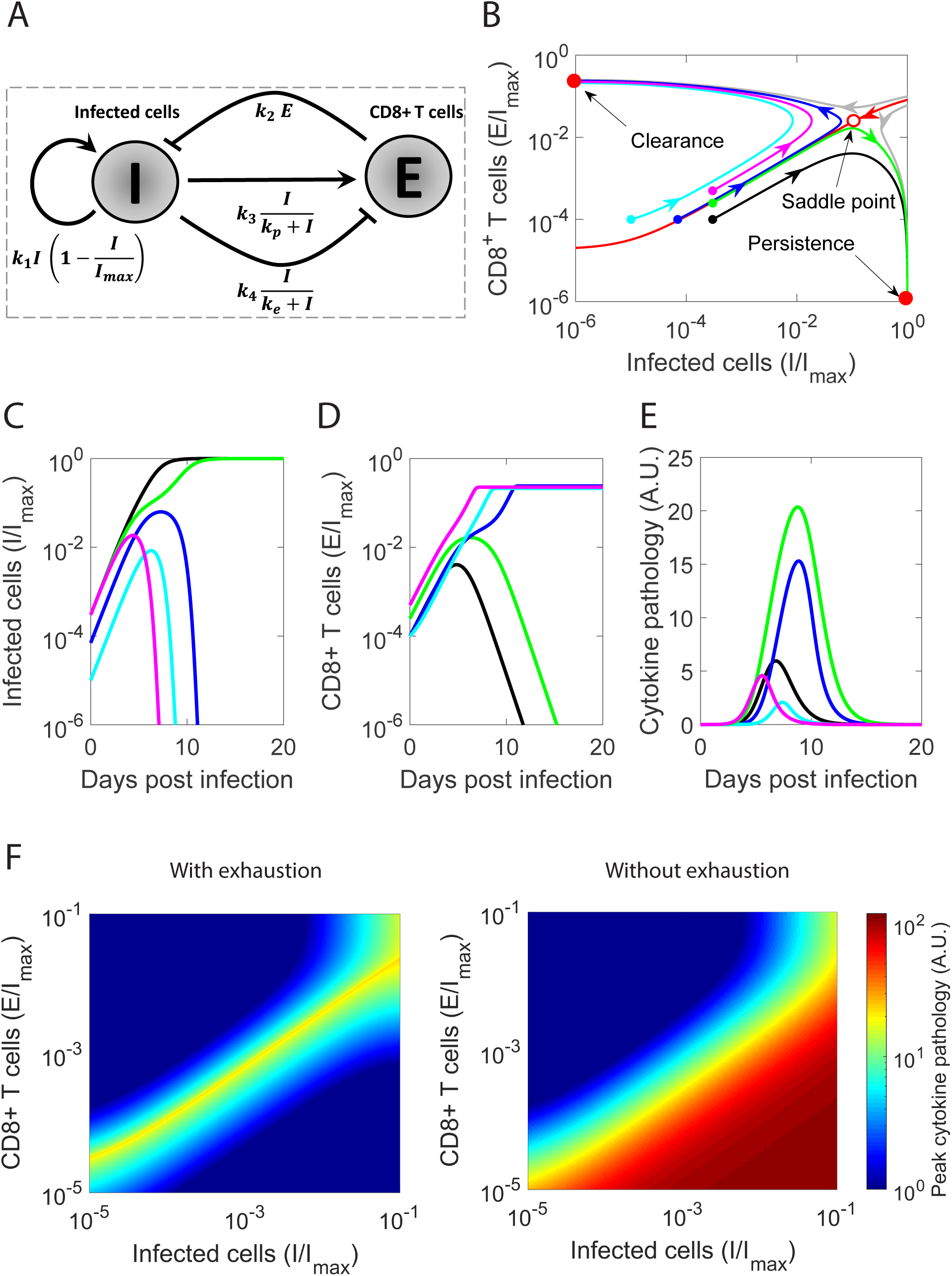
Dynamical motif underlying the outcomes of viral infections (A) Schematic of the dynamical motif depicting the essential interactions between infected cells, *I*, and CD8 T cells, *E*. (B) Phase portrait of the system, where regions leading to the two stable steady states (filled red dots), clearance and persistence, are separated by a basin boundary (red line). The unstable saddle node (open red dot) underlies immunopathology. Also shown are trajectories with increasing initial antigen load (cyan, blue and black) and increasing initial effector pool size (black, green, and magenta) and the corresponding time course of (C) infected cells, (D) effector pool size, and (E) pathology. (F) Heat maps of peak pathology for different initial conditions on the phase portrait with and without exhaustion. Parameters employed are in Supplementary Table S1.

We modelled the growth of infected cells *I* (the source of antigenic-stimulation) using a logistic term with a per capita proliferation rate *k*_1_ and a carrying capacity *I*_*max*_. Infected cells are killed by activated CD8 T cells, *E*, with the second order rate constant *k*_2_. Following previous models (25, 26), we described T cell activation and exhaustion by infected cells using Hill functions with maximal per capita rates *k*_3_ and *k*_4_ and scaling constants *k*_*p*_ and *k*_*e*_, respectively. To ensure that activation happens at lower antigen levels than exhaustion and the latter dominates at high antigen levels, we let *k*_3_ *< k*_4_ and *k*_*p*_ *< k*_*e*_ (26). Thus, as antigen levels rise, they first activate CD8 T cells, whereas continued increase of antigen levels triggers CD8 T cell exhaustion. Our key findings are robust to alternative formulations where exhaustion is triggered by cumulative rather than instantaneous antigen stimulation (20) (Supplementary Text S1; Supplementary Figure S1). We show below that the motif recapitulates the observed outcomes of viral infections.

### Viral clearance and persistence as steady states of the motif

To describe the long-term outcomes of infection, we solved the model equations (Eq. 1) for steady state. The model admits three steady states. The steady states can be derived analytically when the exhaustion rate is far from saturation (*k*_*e*_ ≫ *I*) and can be approximated using the mass action term *k*_4_*IE*. The resulting steady states are: (1) *I* = 0 and *E* ≥ 0; (2) *I* = *I*_*max*_ and *E* = 0; and (3) 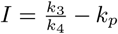 and 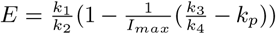. Using linear stability analysis, we found the first two steady states to be stable and the third to be an unstable saddle node (Supplementary Text S2). A phase portrait shows the two stable states separated by a basin boundary on which the saddle node lies (Figure 1B). The first steady state has no antigen and thus marks the clearance of infection. The second, in contrast, has maximal antigen and no effector cells, which indicates suppression of the immune response and persistent infection. (Note that in reality a small population of effector cells will exist in this state, especially if there is continuous immigration from the thymus (26).) The basin boundary (red line in Figure 1B) separates conditions that lead to one outcome from those that lead to the other. Infections initiated in the region above the basin boundary, which marks higher than a threshold CD8 T cells for a given antigen load, lead to viral clearance (Figures 1C,D). In contrast, infections initiated in the region below the basin boundary lead to persistence.

### Immunopathology as a dynamical outcome of the motif

Immunopathology can result from a variety of processes associated with the destruction of virus-infected cells or the release of cytokines resulting in a “cytokine-storm”. Immunopathology is high when the infection spreads to a sizeable proportion of target cells and there is a large population of CD8 T cells kill these cells, directly producing tissue damage or indirectly causing pathology due to cytokine release. We thus modelled immunopathology *P* with the equation *dP/dt* = *αIE* – *d*_*c*_*P*. We found immunopathology to be maximum with intermediate initial antigen load and CD8 T cell counts (Figure 1E). When the initial antigen load is small, CD8 T cells eliminate antigen before it can spread substantially. When the load is large, CD8 T cells CD8+ exhausted before they can kill a sizeable proportion of infected cells. At intermediate antigen loads, the CD8 T cell population is not capable of clearing the infection before it spreads significantly. At the same time, the antigen does not accumulate to extents that can trigger significant CD8 T cell exhaustion. Thus, the antigen spreads and is continuously eliminated by CD8 T cells, which results in sizeable tissue damage.

From a dynamical systems viewpoint, immunopathology results when infection commences close to the basin boundary (Figures 1B,E). When the infection commences away from the boundary, it quickly settles into one of the stable steady states. When it starts close to the boundary, it traverses long trajectories along the boundary, causing a build up of both antigen and CD8 T cell levels and triggering immunopathology before deviating towards one of the stable steady states. Spanning a wide range of initial conditions, we found that conditions close to the basin boundary yielded the maximum immunopathology (Figure 1F).

CD8 T cell exhaustion has been argued to be an evolutionary adaptation to prevent immunopathology (20–22). To test this argument, we repeated our calculations in the absence of exhaustion (*k*_4_ = 0). In contrast to the maximum seen above, we found that pathology now monotonically increased with viral inoculum and effector pool sizes (Figure 1F), demonstrating that exhaustion curtails pathology associated with high viremia and enables host survival. The price, however, is viral persistence.

The motif thus recapitulates the outcomes of viral infections observed. We examined next whether it can describe the roles of key factors known to influence the outcomes.

### Influence of the viral inoculum and initial effector pool sizes

In experiments, infection with low dose LCMV Docile or clone 13 led to clearance, whereas infection with high dose led to persistence in 100% of the mice infected (5) (Figure 2A). With intermediate dose Docile infection, *∼*50% mice cleared the infection and *∼*50% became persistently infected (5), whereas with clone 13 infection, *∼*20% died due to immunopathology and the remaining became persistently infected (6). A small CD8 T cell population at the start of infection, introduced by adoptive transfer, did not prevent persistent LCMV clone 13 infection, whereas a large population led to clearance in the absence of immune escape mutations (27) (Figure 2B). With intermediate effector pool sizes, all the mice infected died due to immunopathology (27).

**Fig. 2.**
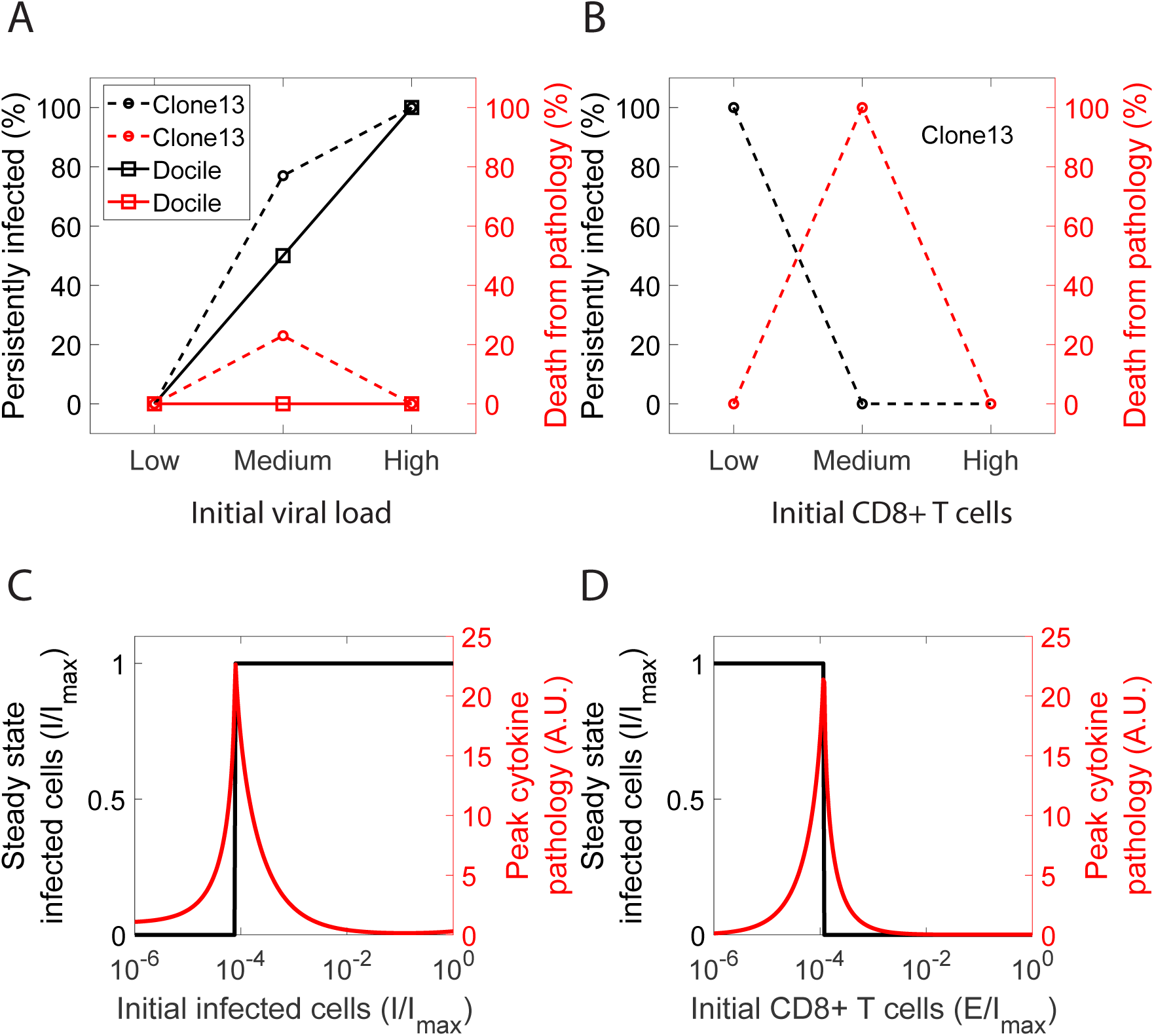
Dependence of outcomes of infection on viral inoculum and initial effector pool sizes. Experimental observations of outcomes with different (A) initial LCMV inoculum sizes (5, 6) and (B) effector pool sizes (27). (C) and (D) Corresponding model predictions of steady state infected cells and peak cytokine pathology. Parameter values are the same as in Figure 1.

In agreement, our model yielded clearance with small viral inocula and persistence with large inocula with fixed initial effector pool sizes (Figure 2C). Immunopathology was maximum with intermediate inocula. This is understood from our predictions above (Figure 1), where as one crosses the basin boundary from small to large viral inocula at a fixed initial CD8 T cell pool size, the outcome switches from clearance to persistence, passing through heightened pathology near the boundary. Similarly, when the initial effector pool size is increased with a constant viral inoculum, the outcome switches from clearance to persistence, with maximum pathology at intermediate sizes (Figure 2D). Host mortality depends on the severity and the criticality of the associated tissue damage. As the initial conditions moved to higher effector cell pool sizes along the basin boundary, we found more severe immunopathology (Figure 1E). Consistent with these predictions, with clone 13 infection, host mortality increased from *∼*20% without to 100% with adoptive transfer of CD8 T cells. With the LCMV Docile infection, the tissue damage was perhaps not critical, resulting in the even split between clearance and persistence, possibly due to stochastic effects and inter-host variations.

The motif thus explains the outcomes realized with different initial viral inocula and effector pool sizes. To describe the influence of other factors, we constructed suitable embellishments of the motif.

### Influence of NK cells

NK cells have been found to influence the outcomes as follows (6). With a normal NK cell response, *∼*25% mice subjected to intermediate dose LCMV clone 13 infection died due to immunopathology, whereas 100% mice administered high dose inocula survived but with persistent infection (Figure 3A inset). In stark contrast, depleting NK cells resulted in 100% survival with intermediate dose and *∼*60% mortality with high dose infection (Figure 3A). NK cells appear inconsequential with low dose infection.

**Fig. 3.**
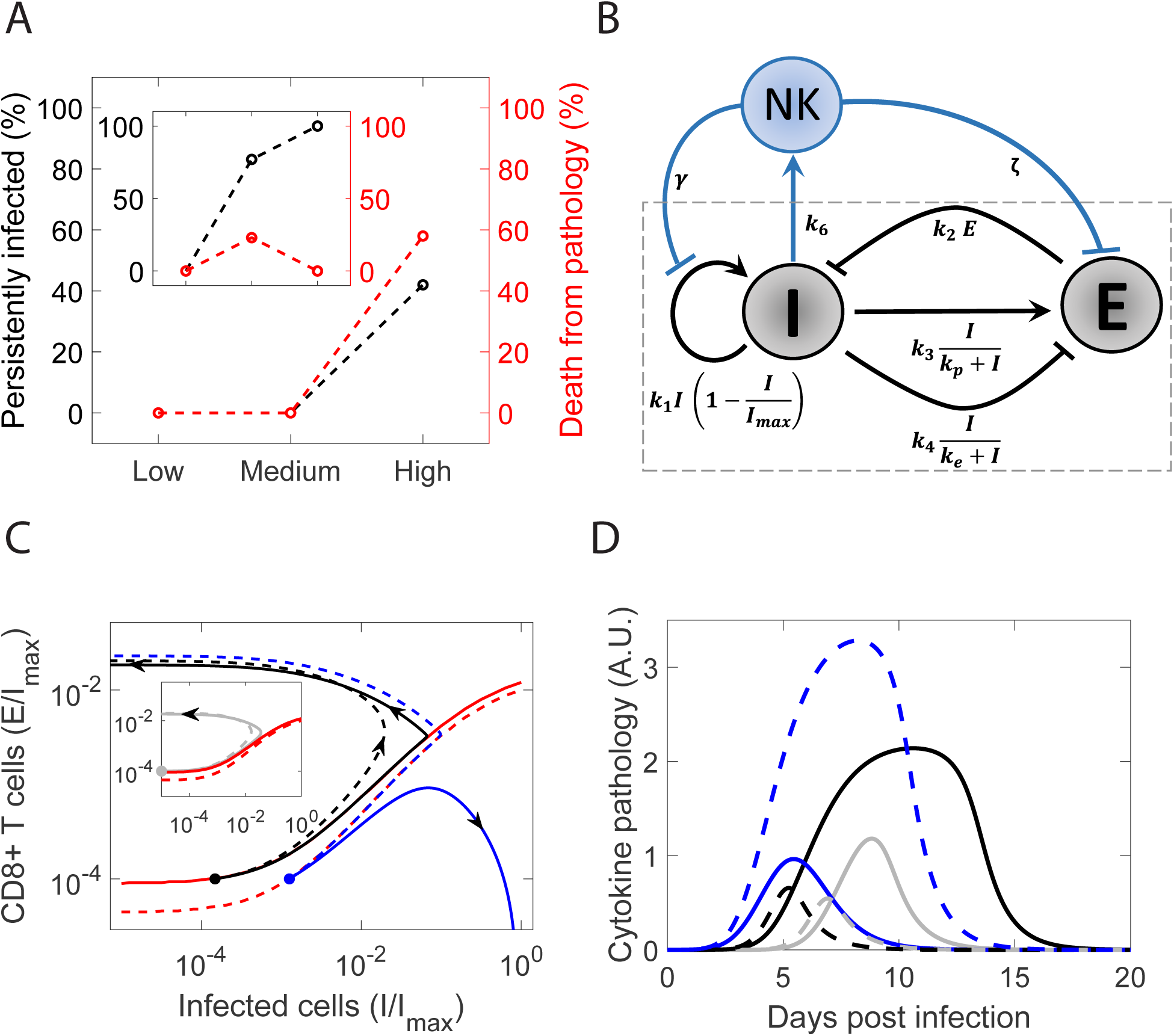
The influence of NK cells. (A) Outcomes observed experimentally in mice infected with different doses of LCMV clone 13 with and without (inset) NK cells (6). (B) Schematic of the motif embellished to incorporate the influence of NK cells. (C) Trajectories corresponding to low (grey; inset), medium (black) and high (blue) dose infection with (solid) and without (dashed) NK cells. The basin boundaries are in red. (D) The corresponding time evolution of cytokine pathology. Parameters employed are in Supplementary Table S1.

NK cells can kill virus-infected cells. However, they can also cytolytically eliminate CD4+ T cells, which can compromise the CD8 T cell response (6) because activation of CD8 T cells typically relies on CD4+ T cell help. We incorporated these effects in the motif by letting NK cells both kill infected cells and suppress the CD8 T cell response (Figure 3B).

Solving the resulting equations (Supplementary Text S3), we found that whereas with normal NK cells, intermediate dose infection led to immunopathology and high dose to persistence and host survival, NK cell depletion reversed the trend with heightened immunopathology with high dose infection (Figure 3C,D). On the phase portrait, depleting NK cells shifted the basin boundary, expanding the basin of attraction of the steady state representing clearance (Figure 3C). This was because the suppression of the effector response by NK cells was relieved with NK cell depletion. (The direct killing of infected cells by NK cells seemed to have less significant an effect on the outcomes.) For the parameters chosen, the intermediate dose infection commenced close to the basin boundary with NK cells, but became removed from the boundary when NK cells were depleted. Conversely, the high dose infection moved closer to the basin boundary when NK cells were depleted (Figure 3C). Thus, immunopathology decreased with intermediate doses and increased with high doses following NK cell depletion (Figure 3D). With low dose infection, NK cells had a minimal influence (Figures 3C-D). Our motif thus recapitulates experimental observations and explains how NK cells function as rheostats (6): they limit the effector response in the dynamical motif to tune the outcome.

### Influence of the innate immune response

The innate immune response is typically mounted early after infection and begins to exert viremic control before the CD8 T cell response is triggered. For instance, IFN, a key innate immune signalling molecule, is secreted following viral infection and triggers the expression of several hundred genes that collectively create an antiviral state in cells, which controls viral replication in infected cells and prevents the infection of uninfected cells (28). The early innate immune response may be viewed, thus, as a way to suppress viremia before the CD8 T cell response is activated. A strong innate immune response would result in a low antigen load at the onset of the CD8 T cell response, which, based on our understanding of the motif above, would facilitate the clearance of infection. To test this hypothesis, we constructed a phenomenological description of the innate immune response, where the innate immune response is triggered by antigen and in turn suppresses the growth of antigen (Figure 4A).

**Fig. 4.**
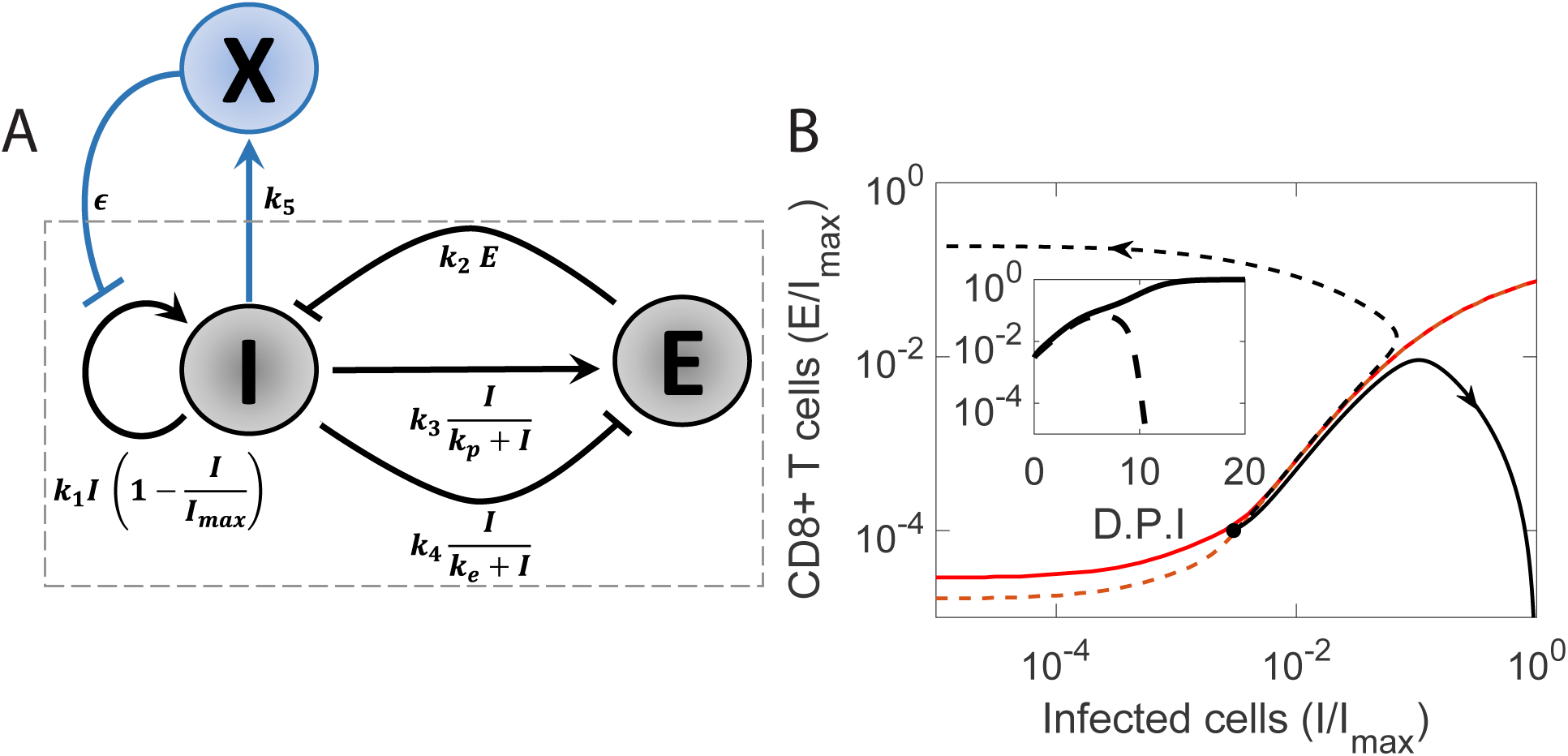
Influence of the innate immune response. (A) Schematic of the motif with the embellishment representing the innate immune response, *X*. (B) Trajectories and time evolution of pathogen load (inset) with a strong (dashed) and weak (solid) innate immune responses. Red lines represent the corresponding basin boundaries. Parameters employed are in Supplementary Table S1.

Solving the resulting model equations (Supplementary Text S4), we found that with all other parameters constant, a strong innate immune response resulted in clearance and a weak response in persistence (Figure 4B inset). Akin to the NK cell response, a strong innate immune response expanded the basin of attraction of the steady state representing clearance (Figure 4B). Unlike the NK cell response, however, which altered the CD8 T cell response, the innate immune response modulated the outcomes by suppressing the antigen load and tilting the balance in favour of effector cells. We considered a realization of the latter phenomenon next, in the spontaneous clearance of HCV infection.

### Spontaneous clearance of HCV infection

Spontaneous clearance of HCV infection has been attributed to strong IFN responses (29, 30). It is also known to require strong CD8 T cell responses (31). To test whether spontaneous clearance is a manifestation of the influence of the innate immune response described above, we constructed a model of HCV dynamics that included the innate immune response. The model combined standard models of HCV kinetics (32, 33) with our motif above (Figure 5A). We solved the resulting equations (Supplementary Text S5) using parameter values representative of HCV infection and examined the influence of the strength of the innate immune response. We found that with a weak innate immune response, the infection became persistent, whereas with a strong innate immune response, the infection was cleared following an acute phase (Figure 5B). We observed further that the rise of viremia during the acute phase was biphasic with a sharp initial rise followed by a slower, more gradual rise to the peak viremia. The arrest of the sharp initial rise was due to the innate immune response. The decline following peak viremia was attributable to the CD8 T cell response (Figure 5B inset). Note that the CD8 T cell response was potentiated by the strong innate immune response, which suppressed viremia and limited exhaustion.

**Fig. 5.**
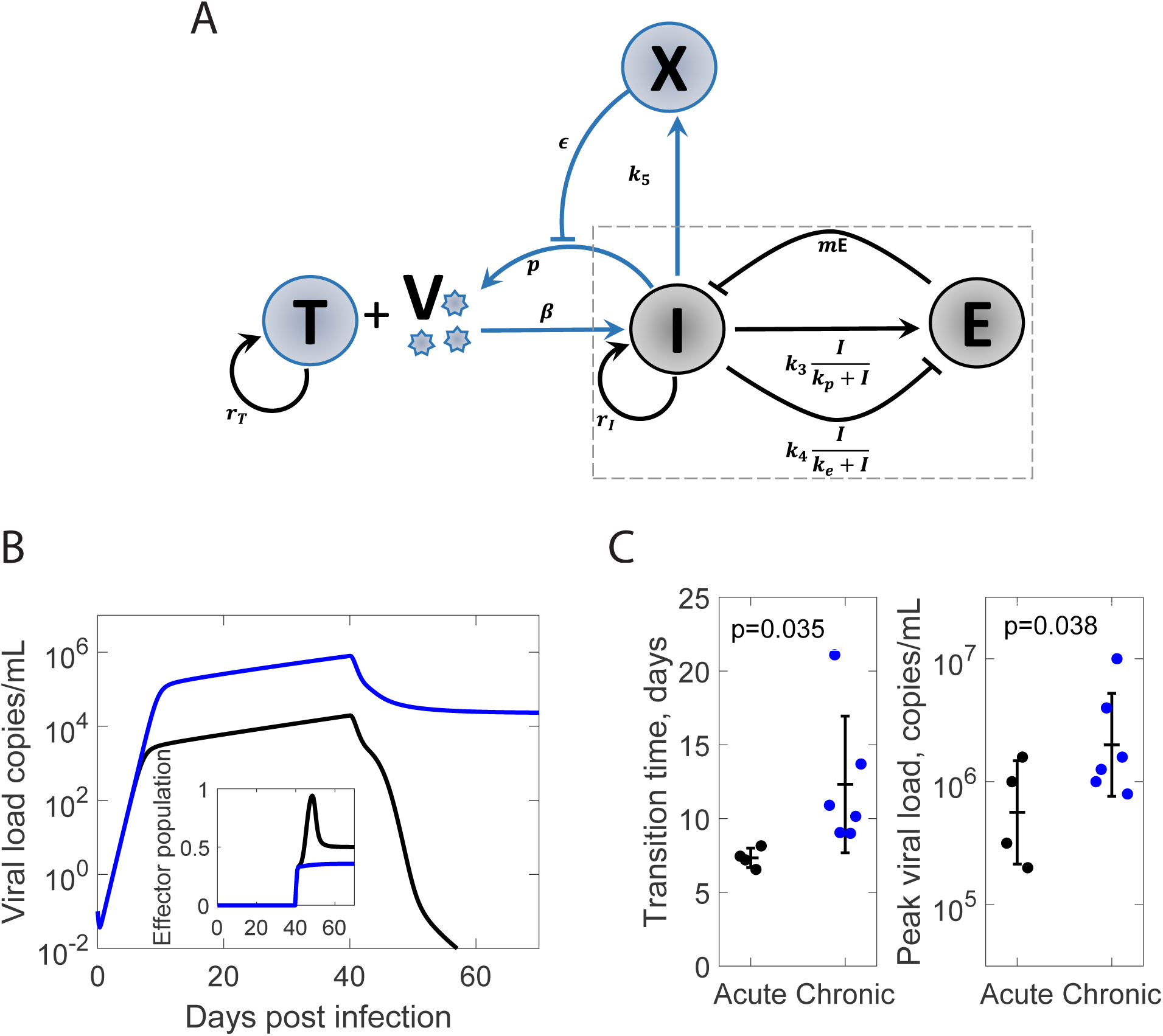
Innate immune response and the outcomes of HCV infection. (A) Schematic of the motif expanded to explicitly include viral dynamics. (B) Time course of viral load and (inset) effector pool size for strong (black) and weak (blue) innate immune responses. Parameters employed are given in Supplementary Table S2. (C) Experimental observations (33, 34) of (left) transition time from first to second phase rise of viremia and (right) peak viral load in chimpanzees infected with HCV resulting in clearance (black) or persistence (blue) of the infection. P values were obtained using one tailed t-tests with equal variance. Parameters employed are in Supplementary Table S2.

Interestingly, the biphasic rise of viremia following infection has been seen in chimpanzees infected with HCV (34). Of the 10 animals challenged, 6 had persistent infection, whereas 4 cleared the infection after the acute phase. We reasoned that the earlier the transition from the first to the second phase rise of viremia, the stronger must be the innate immune response. The resulting peak viremia, which sets the antigen load for the CD8 T cells, would be correspondingly lower, facilitating cure. From previous fits to the viral load data (33), we estimated the transition times and peak viral loads in each animal. We found, remarkably, that the 6 animals that suffered persistent infection had significantly delayed transition times (mean *∼*12 days versus *∼*7 days; P=0.035) and higher peak viral loads (mean *∼*6.3 log10 versus *∼*5.8 log10 copies/ml; P=0.038) than the 4 that cleared the infection (Figure 5D). The data thus support our hypothesis of the innate immune response modulating the outcomes by lowering the antigen load with which the CD8 T cells must contend.

## Discussion

The quest for an understanding of the outcomes of viral infections has led, over the years, to the identification of an array of factors, including the size of the viral inoculum (5), the innate and NK cell responses (6, 30), and host genetic variations (13), that can potentially influence the outcomes. Here, we argue that underlying the effects of these factors is an essential dynamical motif, comprising the key interactions between antigens and CD8 T cells, that governs the outcomes. Using minimal mathematical models of the motif, we recapitulated the different outcomes observed. Broadly, when the CD8 T cell response is strong, the infection is cleared, whereas when the antigen induces significant CD8 T cell exhaustion, long-term persistence results. When the two effects are comparable, substantial spread of infection and associated CD8 T cell killing ensue, causing tissue damage and pathology. Furthermore, using suitable embellishments of the motif, we showed that other factors could influence the outcomes by modulating the interactions in the motif. A new conceptual understanding of the outcomes of viral infections thus emerges. An implication is that, akin to the factors above, interventions that can manipulate the interactions in the motif may present novel routes to clear persistent infections or limit immunopathology.

One intervention strategy, of great interest today, is to use immune checkpoint inhibitors to block inhibitory receptors and reverse CD8 T cell exhaustion. The strategy is being actively explored for treating persistent infections and cancers (35). Our study predicts an advantage to using immune checkpoint inhibitors early in infection. Infections typically commence with small antigen loads and small pools of antigen-specific CD8 T cells. At this stage, immune checkpoint inhibitors may be more readily able to reverse, if not prevent, exhaustion. From a dynamical systems viewpoint, a small increase in effector strength under these circumstances could push the infection across the basin boundary and alter the outcome from persistence to clearance. Once the infection has progressed, however, the level of exhaustion is expected to be higher. Furthermore, the infection is likely to have veered away from the basin boundary towards persistence, requiring a much larger push to drive it to clearance. Our study also sounds a cautionary note. Reinvigorating the immune response in the presence of significant viral antigen may increase pathology. The pathology is likely to be specific to infected tissues and thus distinct from the broad immune-related pathologies seen with checkpoint inhibitors during cancer treatments (36).

As an alternative strategy, antiviral drugs could be used to lower antigen levels, which could alter the outcome from persistence to clearance. Indeed, treatment with the new direct-acting antiviral agents (DAAs) has been found recently to lead to the clearance of HCV infection in many individuals with detectable virus at the end of treatment (37, 38). The mechanisms underlying this post-treatment cure are yet to be established (39–41). Nonetheless, the phenomenon is consistent with our expectation of clearance following a sufficient reduction in antigen load. Post-treatment control has also been seen in a small percentage of HIV infected individuals treated early with antiretroviral therapy (ART) (42). The interpretation, however, is more involved than HCV because of latently infected cells formed by HIV (43), which escape ART and host immune responses and can reignite infection post treatment. One hypothesis is that ART lowers viremia, which reinvigorates exhausted CD8+ T cells and enables control when reignition happens from a small latent cell pool (26). A large latent cell pool can result in large surges of viremia following reactivation, causing exhaustion again and compromising control.

In a series of elegant studies, paradoxical elements, where a stimulus can both activate and suppress its target, have been identified as designs prevalent in biological systems including immune cell circuits (44–46). For instance, the cytokine interleukin 2 can induce the proliferation as well as the death of CD8+ T cells. Such paradoxical pleiotropy has been argued to help maintain homeostatic cell concentrations and robustness in differentiation processes (44, 45). We recognize paradoxical pleiotropy in our motif, where antigen can both activate and suppress CD8+ T cells. An additional feature of our motif is that the target can suppress the stimulus; CD8+ T cells can kill infected cells. Our motif thus represents paradoxical pleiotropy coupled with a double negative feedback loop, the latter also present in many biological networks (47, 48). Furthermore, paradoxical pleiotropy is evident also in the role of NK cells, which can kill infected cells and also compromise effector responses by destroying CD4+ T cells (6, 49). Outcomes of viral infections thus represent a manifestation of paradoxical pleiotropy embedded in a more involved motif including double negative feedback.

Our study has limitations. While our study recapitulates the different outcomes observed, it does not describe the frequencies with which they occur. A distribution of the strength of IFN responses across individuals has been argued to explain the frequency of spontaneous clearance observed with HCV infection (29). Variations in other factors as well as in the strengths of the interactions in our motif, which remain to be quantified, may explain the frequencies observed with other infections. Inter-host variations have been implicated in the differences in the outcomes realized with the same virus (13). In the extreme, a single outcome may be realized, such as with measles virus infection, which is nearly always cleared (50), or HIV infection, which invariably leads to persistence (51). While we have addressed the roles of key factors such as NK cells and innate immune responses, further embellishments of our motif are necessary to describe the influence of other factors. For instance, IFN is known to have both antiviral and immune suppressive effects. Our study has considered the former but not the latter, which is dominant late in infection, and which can be relieved by IFN blockade to achieve clearance of persistent infections (11–13). Finally, whereas in our minimal model CD8+ T cell exhaustion is fully reversible, recent studies have identified subsets of exhausted CD8+ T cells that cannot be reversed, especially with checkpoint inhibitors (52–54). These subsets must be considered in devising accurate strategies to clear persistent infections by manipulating the interactions in our motif.

## ACKNOWLEDGMENTS

The study was supported by the Wellcome Trust/DBT India Alliance Senior Fellowship IA/S/14/1/501307 (NMD) and by NIH U19 AI117891 and R01AI110720 (RA).

