## Supplementary Text, Tables and Figure for "A dynamical motif comprising the interactions between antigens and CD8 T cells may underlie the outcomes of viral infections"

**This PDF file includes:**

Supplementary Text S1 to S5

Supplementary Figure S1

Supplementary Table S1 and S2

References for SI reference citations

### Supporting Information Text

**Supplementary Text S1. Predictions using an alternative model of CD8 T cell exhaustion.** We examined the implications of CD8 T cell exhaustion triggered by cumulative rather than instantaneous antigenic stimulation. Following a previous formalism (1), we wrote:

$$\begin{aligned}\frac{dI}{dt} &= k_1 I\Big(1-\frac{I}{I_{max}}\Big) - k_2 IE \\ \frac{dE}{dt} &= k_3\frac{IE}{k_p+I} - k_4\frac{QE}{k_e+Q} \\ \frac{dQ}{dt} &= q_S\Big(\frac{I}{\phi+I} - Q\Big)\end{aligned} \qquad [S1]$$

Here, $Q$ represents the cumulative stimulation level, which increases with antigen load in a saturable manner with half-maximal constant $\phi$ and decays with the rate constant $q_S$. $Q$ then determines the level of exhaustion, as opposed to the antigen load in Eq. 1 in the main text, and suppresses the effector response. The resulting motif is illustrated in Figure S1A. Solving the equations above yielded clearance with low viral inoculum sizes and/or high initial effector pool size, immunopathology with intermediate sizes, and persistence with high viral inoculum sizes and/or low initial effector pool sizes (Figure S1). The specific forms used to defining exhaustion thus did not alter our key conclusions.

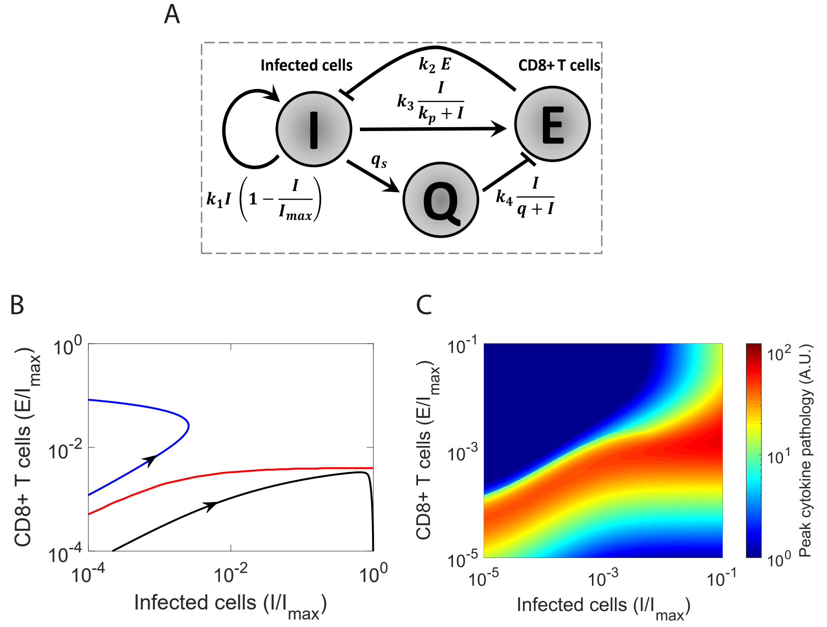


Figure S1: Dynamical motif with exhaustion induced by cumulative antigen stimulation. (A) A schematic of the motif described in Supplementary Text S1. (B) Phase portrait showing the basin boundary (red line) separating initial conditions that lead to clearance (above red line) and persistence (below red line). Blue and black lines show projections of trajectories leading to the two outcomes. (Note that the system is now high dimensional and the projections can thus cross the basin boundary.) (C) Cytokine pathology associated with a range of initial conditions illustrating maximum pathology near the basin boundary. Parameter values employed are as follows: $k_e = 0.8$; $q_s = 0.08$/day; and $\phi = 1000$ cells (1). The other parameters are the same as in Figure 1.

**Supplementary Text S2. Linear stability analysis.** The system in Eq. 1 in the main text with the exhaustion term away from saturation has 3 steady states. To determine the stability of these steady states, we performed linear stabililty analysis. The Jacobian matrix of the system is

$$\mathrm{A} = \begin{bmatrix} k_1 - \frac{2k_1 I}{I_{max}} - k_2 E & -k_2 I \\ \frac{k_3 k_p E}{(I+k_p)^2} - k_4 E & \frac{k_3 I}{I+k_p} - k_4 I \end{bmatrix} \qquad [S2]$$

For the steady state representing clearance ($I = 0$ and $E \geq 0$), the Jacobian becomes

$$\mathrm{A} = \begin{bmatrix} k_1 - k_2 E & 0 \\ \frac{k_3 E}{k_p} - k_4 E & 0 \end{bmatrix} \qquad [S3]$$

Thus, the $trace(A) = k_1 - k_2 E$ and $det(A) = 0$, which would yield a line of stable fixed points as long as $E \geq \frac{k_1}{k_2}$.

For the steady state representing persistence ($I = I_{max}$ and $E = 0$), the Jacobian becomes

$$\mathrm{A} = \begin{bmatrix} -k_1 & -k_2 I_{max} \\ 0 & \frac{k_3 I_{max}}{I_{max}+k_p} - k_4 I_{max} \end{bmatrix} \qquad [S4]$$

which has both eigenvalues real and negative if $\frac{k_3}{k_4} - k_p < I_{max}$. This steady state is thus stable as long as the latter inequality holds.

For the third steady state ($I = \frac{k_3}{k_4} - k_p$ and $E = \frac{k_1}{k_2}(1 - \frac{1}{I_{max}}(\frac{k_3}{k_4} - k_p))$), we can show that the signs of the elements of the Jacobian are

$$\mathrm{A} = \begin{bmatrix} + & - \\ - & 0 \end{bmatrix} \qquad \text{[S5]}$$

Thus, $det(A) < 0$, so that the eigenvalues have opposite signs. The steady state is therefore a saddle point.

**Supplementary Text S3: Mathematical model of the motif with NK cells.** The following equations describe the behavior of the motif with NK cells (Figure 3):

$$\begin{aligned} \frac{dI}{dt} &= k_1 I\big(1 - \frac{I}{I_{max}}\big) - k_2 IE - \gamma IN \\ \frac{dE}{dt} &= k_3\big(1 - \zeta\frac{N}{\phi_E + N}\big)\frac{IE}{k_p + I} - k_4\frac{IE}{k_e + I} \\ \frac{dN}{dt} &= k_6\big(\frac{I}{\phi_N + I} - N\big) \end{aligned} \qquad \text{[S6]}$$

Here, we let the NK cell response, $N$, rise in proportion to the antigen load, $I$, in a saturable manner with the scaling constant $\phi_N$ and decay with the rate constant $k_6$. Further, we let the NK cells eliminate infected cells at the rate $\gamma$ and suppress the CD8 T cell response by the maximal extent $\zeta$. The remaining terms are similar to those in Eq. 1 in the main text.

**Supplementary Text S4: Mathematical model of the motif with the innate immune response.** The following equations describe the behavior of the motif with the innate immune response (Figure 4):

$$\begin{aligned} \frac{dI}{dt} &= k_1\big(1 - \epsilon\frac{X}{\phi_X + X}\big)I\big(1 - \frac{I}{I_{max}}\big) - k_2 IE \\ \frac{dE}{dt} &= k_3\frac{IE}{k_p + I} - k_4\frac{IE}{k_e + I} \\ \frac{dX}{dt} &= k_5\big(\frac{I}{\phi_I + I} - X\big) \end{aligned} \qquad \text{[S7]}$$

Here, $X$ is a coarse-grained, normalized variable quantifying the innate immune response. $X$ rises in proportion to the antigen load, $I$, in a saturable manner with the scaling constant $\phi_I$ and decays with the rate constant $k_5$. We assumed that $X$ restricts the spread of infection with the maximal efficacy $\epsilon$ and the scaling constant $\phi_X$. $\phi_I$ sets the scale of the antigen load that triggers the innate immune response. Large values of $\phi_I$ imply weak immune responses. The remaining terms are similar to those in Eq. 1 in the main text.

**Supplementary Text S5: Model of HCV dynamics with innate immune response.** We constructed a model of HCV dynamics that included our framework above of the innate immune response (Figure 3A) in standard models of HCV kinetics (2, 3) (see Figure 4A). The model equations are as follows:

$$\begin{aligned} \frac{dT}{dt} &= s + r_T T\big(1 - \frac{T+I}{T_{max}}\big) - d_T T - \beta VT + f_2 mIE \\ \frac{dI}{dt} &= \beta VT + r_I I\big(1 - \frac{T+I}{T_{max}}\big) - \delta I - m(f_1 + f_2)IE \\ \frac{dV}{dt} &= p\big(1 - \epsilon\frac{X}{\phi_X + X}\big) - cV \\ \frac{dX}{dt} &= k_5\big(\frac{I^n}{\phi_I^n + I^n} - X\big) \\ \frac{dE}{dt} &= \lambda\mathcal{H}(\tau_d) + k_3\frac{IE}{k_p + I} - k_4\frac{IE}{k_e + I} - \mu E \end{aligned} \qquad \text{[S8]}$$

Here, $T$ represents the population of target, uninfected hepatocytes; $I$, infected hepatocytes; $V$, virus; $X$, the innate immune response; and $E$, CD8 T cells. Target cells are produced at the rate $s$, proliferate at the per capita rate $r_T$, die at the per capita rate $d_T$, and are infected by virions at the rate $\beta VT$, to yield infected cells. The latter cells proliferate at the per capita rate $r_I$, and die at the per capita rate $\delta$. Total cell proliferation is restricted by a logistic term with carrying capacity $T_{max}$. Virus is produced at the rate $p$ per infected cell and cleared at the per capita rate $c$. These latter steps represent details of antigenic growth that we had subsumed into the proliferation of infected cells in our dynamical motif above (Figure 1A).

We let the innate immune response follow the form above (Eq. S7). Additionally, we allowed the sensitivity of the response to infected cells to be modulated by the Hill co-efficient $n$. The response suppresses viral production (2), which we assumed to occur with the maximal efficacy $\epsilon$. For CD8 T cell dynamics, we followed earlier models of activation and exhaustion (1, 4) together with a delay in the induction of the response (5–7), characterized using $\tau_d$ (3). $\mathcal{H}$ represents the Heaviside function such that $\mathcal{H}(x) = 0$ if $x < 0$ and $\mathcal{H}(x) = 1$ if $x \geq 0$. CD8 T cells are thus produced at the rate $\lambda$ after a delay of $\tau_d$ following the onset of infection and die at the per capita rate $\mu$. Their effect on infected cells is both cytolytic and non-cytolytic (3). We represented the two effects using the terms $f_1$ and $f_2$, respectively. Thus, infected cells are destroyed at the rate $mf_1IE$ and recover and become target cells at the rate $mf_2IE$.

**Table S1. Parameter estimates. The parameter values employed in our model calculations are listed along with ranges available in the literature. Where no estimates were available, we have assumed parameters values to ensure that the key qualitative features of the system are maintained.**

| Parameter | Meaning | Value | Range or value from literature | Source |
|---|---|---|---|---|
| | | **Figures 1 & 2** | | |
| $k_1$ | Infection spread rate | 1.3/day | 3-4/day | (8, 9) |
| $I_{max}$ | Maximum number of infected cells | $10^6$ cells | $10^6$ cells | (1) |
| $k_2$ | Killing rate of infected cells by CD8 T cells | $5 \times 10^{-5}$/cells/day | $\sim 10^{-6} - 10^{-4}$ /cells/day | (8, 9) |
| $k_3$ | Antigen driven CD8 T cell proliferation rate | 1/day | $\sim$1/day | (1, 4, 10) |
| $k_4$ | Antigen driven CD8 T cell suppression rate | 3/day | 2-4/day | (1, 4, 10) |
| $k_p$ | Antigen driven proliferation threshold | 10 cells | $10^{-1} - 10^3$ cells | (1, 4) |
| $k_e$ | Antigen driven suppression threshold | $2 \times 10^5$ cells | $5 - 2.7 \times 10^4$ cells | (4, 10) |
| $\alpha$ | Rate of growth of cytokine pathology | $2 \times 10^{-8}$ /cells$^2$/day | | |
| $d_c$ | Rate of loss of cytokine pathology | $1$/day | | |
| | | **Figure 3** | | |
| $k_1$ | Infection spread rate | 1.5/day | 3-4/day | (8, 9) |
| $I_{max}$ | Maximum number of infected cells | $10^6$ cells | $10^6$ cells | (1) |
| $k_2$ | Killing rate of infected cells by CD8 T cells | $1 \times 10^{-4}$/cells/day | $10^{-6} - 10^{-4}$ | (8, 9) |
| $k_3$ | Antigen driven CD8 T cell proliferation rate | 1.55/day | $\sim$1/day | (1, 4, 10) |
| $k_4$ | Antigen driven CD8 T cell suppression rate | 3/day | 2-4/day | (1, 4, 10) |
| $k_p$ | Antigen driven proliferation threshold | 1000 cells | $10^{-1} - 10^3$ cells | (1, 4) |
| $k_e$ | Antigen driven suppression threshold | $1 \times 10^5$ cells | $5 - 2.7 \times 10^4$ cells | (4, 10) |
| $k_6$ | NK cell activation rate | 5/day | | |
| | NK cell activation rate with NK cell depletion | 0/day | | |
| $\phi_N$ | NK cell efficacy threshold | 100 cells | | |
| $\zeta$ | Efficacy of NK cell suppression of CD8 T cell proliferation | 0.3 | | |
| $\phi_E$ | NK cell activation threshold | 0.5 cells | | |
| $\gamma$ | Rate of elimination of infected cells by NK cells | 0.1 cells/day | | |
| $\alpha$ | Rate of growth of cytokine pathology | $2 \times 10^{-8}$ /cells$^2$/day | | |
| $d_c$ | Rate of loss of cytokine pathology | $1$/day | | |
| | | **Figure 4** | | |
| $k_1$ | Infection spread rate | 1.5/day | 3-4/day | (8, 9) |
| $I_{max}$ | Maximum number of infected cells | $10^6$ cells | $10^6$ cells | (1) |
| $k_2$ | Killing rate of infected cells by CD8 T cells | $4 \times 10^{-5}$/cells/day | $10^{-6} - 10^{-4}$ | (8, 9) |
| $k_3$ | Antigen driven CD8 T cell proliferation rate | 1.55/day | $\sim$1/day | (1, 4, 10) |
| $k_4$ | Antigen driven CD8 T cell suppression rate | 3/day | 2-4/day | (1, 4, 10) |
| $k_p$ | Antigen driven proliferation threshold | 1000 cells | $10^{-1} - 10^3$ cells | (1, 4) |
| $k_e$ | Antigen driven suppression threshold | $1 \times 10^5$ cells | $5 - 2.7 \times 10^4$ cells | (4, 10) |
| $k_5$ | Innate immune response activation rate | 10/day | | |
| $\phi_I$ | Innate immune response activation threshold (strong response) | 2000 cells | | |
| | Innate immune response activation threshold (weak response) | 4000 cells | | |
| $\zeta$ | Efficacy of innate immune response suppression of viral replication | 0.95 | | |
| $\phi_X$ | Efficacy threshold for innate immune response | 0.8 cells | | |
| $\alpha$ | Rate of growth of cytokine pathology | $2 \times 10^{-8}$ /cells$^2$/day | | |
| $d_c$ | Rate of loss of cytokine pathology | $1$/day | | |

**Table S2. Parameter values used for our calculations of HCV dynamics (Figure 5).**

| Parameters | Meaning | Value | Source |
|---|---|---|---|
| $s$ | Target cell production rate | 61200 cells/mL/day | (3) |
| $r_T$ | Target cell proliferation rate | 0.53/day | (3) |
| $T_{max}$ | Carrying capacity of liver | $7\times10^{7}$ cells/mL | (3) |
| $d_T$ | Target cell death rate | 0.0034 /day | (3) |
| $\beta$ | Infection rate constant | $1\times10^{-8}$ mL/cells/day | (3) |
| $r_I$ | Infected cell proliferation rate | 0.53/day | (3) |
| $\delta$ | Death rate of infected cells | $1.5\times d_T$ | (3) |
| $m$ | Effector strength of CTL response | 5 mL/cells/day | |
| $f_1$ | Cytolytic fraction of CTL response | 0.3 | (3) |
| $f_2$ | Non-cytolytic fraction of CTL response | 0.7 | (3) |
| $p$ | Virus production rate | 20 virions/cell/day | (3) |
| $\epsilon$ | Efficacy of innate immune response in blocking viral production | 0.98 | (3) |
| $k_5$ | Innate immune response activation rate | 5/day | |
| $\phi_I$ | Innate immune response activation threshold (strong response) | $10^3$ cells/ml | |
| | Innate immune response activation threshold (weak response) | $5\times10^4$ cells/ml | |
| $\lambda$ | $CD8^+$ T cell recruitment rate | 1 cells/mL/day | (4) |
| $k_3$ | Antigen driven CD8 T cell proliferation rate | 1/day | (4) |
| $k_4$ | Antigen driven CD8 T cell suppression rate | 2/day | (4) |
| $k_p$ | Antigen driven CD8 T cell proliferation threshold | 0.1 cells/mL | (4) |
| $k_e$ | Antigen driven CD8 T cell suppression threshold | 1000 cells/mL | |
| $\mu$ | Death rate of $CD8^+$ T cells | 2/day | (4) |
